# Antimicrobial Profile from Frog Skin Peptides

**DOI:** 10.1101/2024.07.03.600345

**Authors:** Yan Chi, Yu Zhang, Xuejiang Wang, Feng Li, Zhikai Zhang

## Abstract

This study evaluates the antimicrobial efficacy of frog skin-derived peptides Magainin 1, Magainin 2, and Bombesin against *Bacillus subtilis, Escherichia coli,* and *Trichoderma harzianum*. Experimental setups involved uniform inoculation of the microorganisms on 20 mL agar plates, treated with varying volumes (1.5 µL, 5 µL, and 10 µL) of each peptide (10 mg/mL). For *Bacillus subtilis*, Magainin 2, and Bombesin demonstrated significant antibacterial activity, with inhibition zones increasing in size proportionally to the peptide volumes. Magainin 2 showed the highest efficacy, surpassing ampicillin at higher volumes. In *Escherichia coli*, similar dose-dependent antibacterial effects were observed, with Magainin 2 again showing superior performance, matching or exceeding the inhibition zones of ampicillin. Notably, Magainin 2, and Bombesin exhibited antifungal activity against *Trichoderma harzianum* with Amphotericin resistance. These antibacterial peptides show inhibitory activity on fungi, gram-positive higher than gram-negative bacteria. These findings highlight the potential of Magainin 2, and Bombesin as antimicrobial agents except of Magainin 1.

## Introduction

The increasing prevalence of antibiotic-resistant bacteria has become a significant global health concern, necessitating the development of new and effective antimicrobial agents. Natural antimicrobial peptides (AMPs) derived from frog skin have garnered attention due to their broad-spectrum activity and potential to overcome resistance mechanisms^1,2^. Among these peptides, Magainin 1^3^, Magainin 2^4^, and Bombesin have shown promising antimicrobial properties^4^. This study aims to evaluate the efficacy of these frog skin-derived peptides against common bacterial pathogens, *Bacillus subtilis* and *Escherichia coli*, as well as the mold *Trichoderma harzianum*.

*Bacillus subtilis* is a Gram-positive bacterium widely used in research due to its well-characterized genetic and physiological properties. It serves as a model organism for studying bacterial growth, differentiation, and pathogenesis^5^. *Escherichia coli*, a Gram-negative bacterium, is another extensively studied microorganism, often used as an indicator of antimicrobial efficacy due to its relevance in various infections^6^. Evaluating the effectiveness of AMPs against these two bacterial species can provide insights into their potential as broad-spectrum antibacterial agents. *Trichoderma harzianum* is a mold known for its role in biocontrol and its resistance to many conventional antifungal agents^7,8^. The ability to inhibit or kill this mold is particularly important in agricultural and environmental applications. However, the effectiveness of AMPs against fungi, particularly molds like *T. harzianum*, remains underexplored. This study seeks to fill this gap by assessing the antifungal activity of Magainin 1, Magainin 2, and Bombesin.

The increasing prevalence of antibiotic-resistant bacteria has become a significant global health concern, necessitating the development of new and effective antimicrobial agents^9,10^. Previous research has demonstrated that AMPs exert their antimicrobial effects by disrupting microbial cell membranes, leading to cell lysis and death.^11,12^. This mechanism differs from traditional antibiotics, which often target specific bacterial processes, thereby reducing the likelihood of resistance development. Given their unique mode of action and broad-spectrum activity, AMPs such as Magainin 1^3^, Magainin 2^13,14^, and Bombesin^15^ hold promise as alternative or adjunctive therapies to combat antibiotic-resistant infections.

In this study, we conducted a series of experiments to evaluate the antimicrobial efficacy of these peptides against Bacillus subtilis, Escherichia coli, and Trichoderma harzianum. We employed a range of peptide concentrations to determine the dose-dependent effects and included ampicillin as a standard antibiotic control. By systematically measuring the inhibition zones, we aimed to provide a comprehensive assessment of the antibacterial and antifungal properties of these frog skin-derived peptides, thereby contributing to the development of new antimicrobial strategies.

## Materials and methods

### Materials

To assess the antimicrobial efficacy of frog skin-derived peptides Magainin 1, Magainin 2, and Bombesin, a series of experiments were conducted against *Bacillus subtilis, Escherichia coli*, and *Trichoderma harzianum*. For Bacillus subtilis, agar plates were uniformly inoculated with the bacteria and treated with three different volumes (1.5 µL, 5 µL, and 10 µL) of each peptide. The clear inhibition zones around the application points were measured to evaluate antibacterial activity. For *Escherichia coli*, a similar setup was employed using petri dishes uniformly inoculated with the bacteria. Different volumes (1.5 µL, 5 µL, and 10 µL) of each peptide were applied, and control groups treated with ampicillin were included for comparison. The formation of clear inhibition zones was observed and measured to assess dose-dependent antibacterial effects. For *Trichoderma harzianum*, petri dishes were uniformly inoculated with the mold and treated with the same volumes (1.5 µL, 5 µL, and 10 µL) of each peptide, with ampicillin-treated control groups included. The plates were incubated under standard conditions suitable for mold growth, and the presence or absence of inhibition zones was documented to evaluate antifungal efficacy. This comprehensive setup allowed for the assessment of the antimicrobial properties of the peptides across different microorganisms and their comparison with a standard antibiotic.

### HPLC analysis

The HPLC analysis of AMP was conducted using a Symmetrix ODS-R column (4.6 x 250 mm, 5 μm) under a gradient elution program. Solvent A consisted of 0.1% trifluoroacetic acid in 100% acetonitrile, while Solvent B was composed of 0.1% trifluoroacetic acid in 100% water. The gradient elution started with 10% Solvent A and 90% Solvent B at 0.0 minutes. The proportion of Solvent A was increased linearly to 100% over 25.0 minutes, maintaining this composition until 25.1 minutes, and the run was terminated at 30.0 minutes. The flow rate was set at 1.0 ml/min, and the detection wavelength was 220 nm. A sample volume of 10 μl was injected for analysis.

The chromatographic conditions provided efficient separation and detection of AMP. The initial high percentage of Solvent B ensured good solubility and initial retention of AMP on the column. As the gradient progressed, increasing Solvent A facilitated the elution of AMP from the column, which was detected at 220 nm. The smooth transition from 10% to 100% Solvent A over 25 minutes allowed for a gradual elution, minimizing potential peak broadening and ensuring a sharp, well-defined peak for AMP. The choice of trifluoroacetic acid in both solvents likely helped improve peak shape and resolution by minimizing tailing effects, making this method effective for analyzing AMP in various samples.

### ESI MASS analysis

The ESI mass spectrometry analysis of AMP was conducted on May 21, 2024, using an ESI probe with a bias of +4.5 kV. The analysis was performed by Huang, with a nebulizer gas flow rate of 1.5 L/min and a detector set to 1.2 kV. The sample analyzed was identified as PGlu-QM-13-NH2. Critical settings for the analysis included a CDL voltage of −20.0 V, a total flow rate of 0.2 ml/min, and a CDL temperature of 250°C. The solvent composition was balanced at 50% water and 50% acetonitrile (ACN), ensuring optimal ionization conditions. The block temperature was maintained at 400°C to facilitate efficient desolvation of the sample.

The ESI probe parameters and the controlled environmental conditions ensured the stability and precision of the mass spectrometric measurements. The combination of a +4.5 kV probe bias and a detector voltage of 1.2 kV allowed for the sensitive detection of AMP ions. The specific sample preparation and solvent composition facilitated the effective ionization of AMP, leading to accurate mass/charge (m/z) ratio readings. The high CDL and block temperatures further improved the desolvation process, minimizing noise and enhancing the clarity of the spectra. This robust analytical setup enabled the reliable identification and quantification of AMP in the PGlu-QM-13-NH2 sample.

### The Antimicrobial Efficacy of Magainin 1, Magainin 2, and Bombesin on Bacillus subtilis

To assess the antimicrobial efficacy of frog skin-derived peptides Magainin 1, Magainin 2, and Bombesin against Bacillus subtilis, an experimental setup was employed involving uniform inoculation of Bacillus subtilis on agar plates. Each plate was treated with three different volumes (1.5 µL, 5 µL, and 10 µL) of the peptides. The clear inhibition zones around the application points were measured to evaluate the peptides’ antibacterial activity, demonstrating a dose-response relationship and highlighting their potential as effective antimicrobial agents.

### The Antimicrobial Efficacy of Magainin 1, Magainin 2, and Bombesin on *Escherichia coli*

To evaluate the antimicrobial efficacy of frog skin-derived peptides Magainin 1, Magainin 2, and Bombesin against Escherichia coli, an experimental setup was employed using petri dishes uniformly inoculated with E. coli. Different volumes (1.5 µL, 5 µL, and 10 µL) of each peptide were applied to the dishes. Control groups treated with ampicillin were also included for comparison. Each petri dish was observed for the formation of clear inhibition zones, which indicate areas where bacterial growth was inhibited by the peptides. The diameter of these zones was measured to assess the effectiveness of each peptide at the different concentrations. This setup allowed for the assessment of dose-dependent antibacterial effects and comparison with a standard antibiotic control.

### The Antimicrobial Efficacy of Magainin 1, Magainin 2, and Bombesin on *Trichoderma harzianum*

To evaluate the antifungal efficacy of frog skin-derived peptides Magainin 1, Magainin 2, and Bombesin against Trichoderma harzianum, an experimental setup was employed using petri dishes uniformly inoculated with T. harzianum. Different volumes (1.5 µL, 5 µL, and 10 µL) of each peptide were applied to the dishes. Control groups treated with ampicillin were included for comparison. Each petri dish was observed for the formation of clear inhibition zones, which would indicate areas where fungal growth was inhibited by the peptides. The plates were incubated under standard conditions suitable for mold growth, and the presence or absence of inhibition zones was documented to assess the effectiveness of each peptide and the antibiotic control in inhibiting the growth of T. harzianum. This setup allowed for the assessment of potential antifungal effects of the peptides and their comparison with a known antibiotic.

## Results

### ESI Mass Spectrometry Analysis of Magainin 1, Magainin 2, and Bombesin

Magainin 1 Analysis: The ESI mass spectrum for Magainin 1 (top panel) displays several significant peaks corresponding to various charge states of the molecule. The most prominent peak is observed at m/z 603.25, corresponding to the [M+4H]4+ ion. Other notable peaks include those at m/z 804.20 ([M+3H]3+) and m/z 1205.80 ([M+2H]2+). These observed peaks indicate the ionization pattern of Magainin 1. To calculate the molecular weight (MW) of Magainin 1, we use the most prominent peak [M+4H]4+. The calculation is as follows:

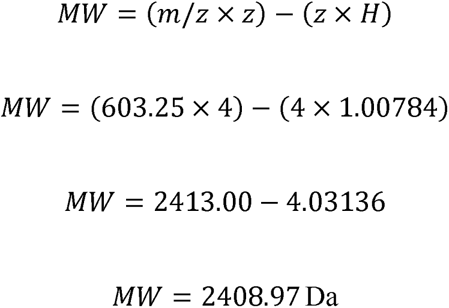

This calculated molecular weight of approximately 2409 Da is consistent with the theoretical molecular weight of Magainin 1.

Magainin 2 Analysis: The ESI mass spectrum for Magainin 2 (middle panel) shows prominent peaks at m/z 617.80 ([M+4H]4+), m/z 823.25 ([M+3H]3+), and m/z 1234.30 ([M+2H]2+). These peaks indicate the ionization states of Magainin 2. Using the [M+4H]4+ peak for molecular weight calculation:

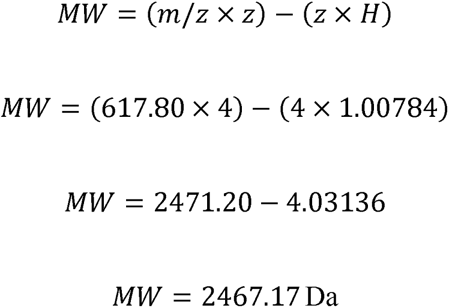

The calculated molecular weight of approximately 2467 Da aligns with the theoretical molecular weight of Magainin 2, confirming the accuracy of the ESI mass spectrometric analysis.

Bombesin Analysis: The ESI mass spectrum for Bombesin (bottom panel) reveals peaks at m/z 821.45 ([M+4H]4+), m/z 1095.80 ([M+3H]3+), and m/z 1645.45 ([M+2H]2+). To calculate the molecular weight of Bombesin, we use the [M+4H]4+ peak:

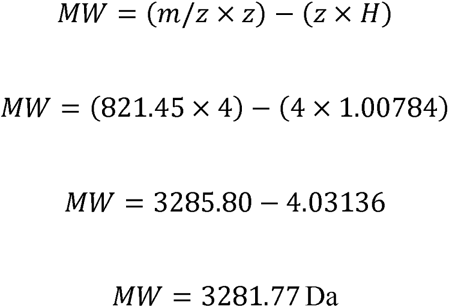

The calculated molecular weight of approximately 3282 Da is consistent with the theoretical molecular weight of Bombesin. This confirms the successful identification and characterization of Bombesin via ESI mass spectrometry.

These results validate the ESI mass spectrometric method used, as the calculated molecular weights for Magainin 1, Magainin 2, and Bombesin closely match their respective theoretical molecular weights, demonstrating the precision and reliability of this analytical technique.

### HPLC Analysis of Magainin 1, Magainin 2, and Bombesin

The HPLC chromatograms presented in Figure 1 reveal the elution profiles and purity of Magainin 1, Magainin 2, and Bombesin under the specified analytical conditions. The separation was conducted using a mobile phase consisting of 50% water and 50% acetonitrile at a flow rate of 0.2 ml/min. The column temperature was maintained at 250°C, with a block temperature of 400°C. Detection was achieved with a detector voltage of 1.2 kV, which ensured high sensitivity and precise identification of the peaks corresponding to each peptide.

**Figure 1:**
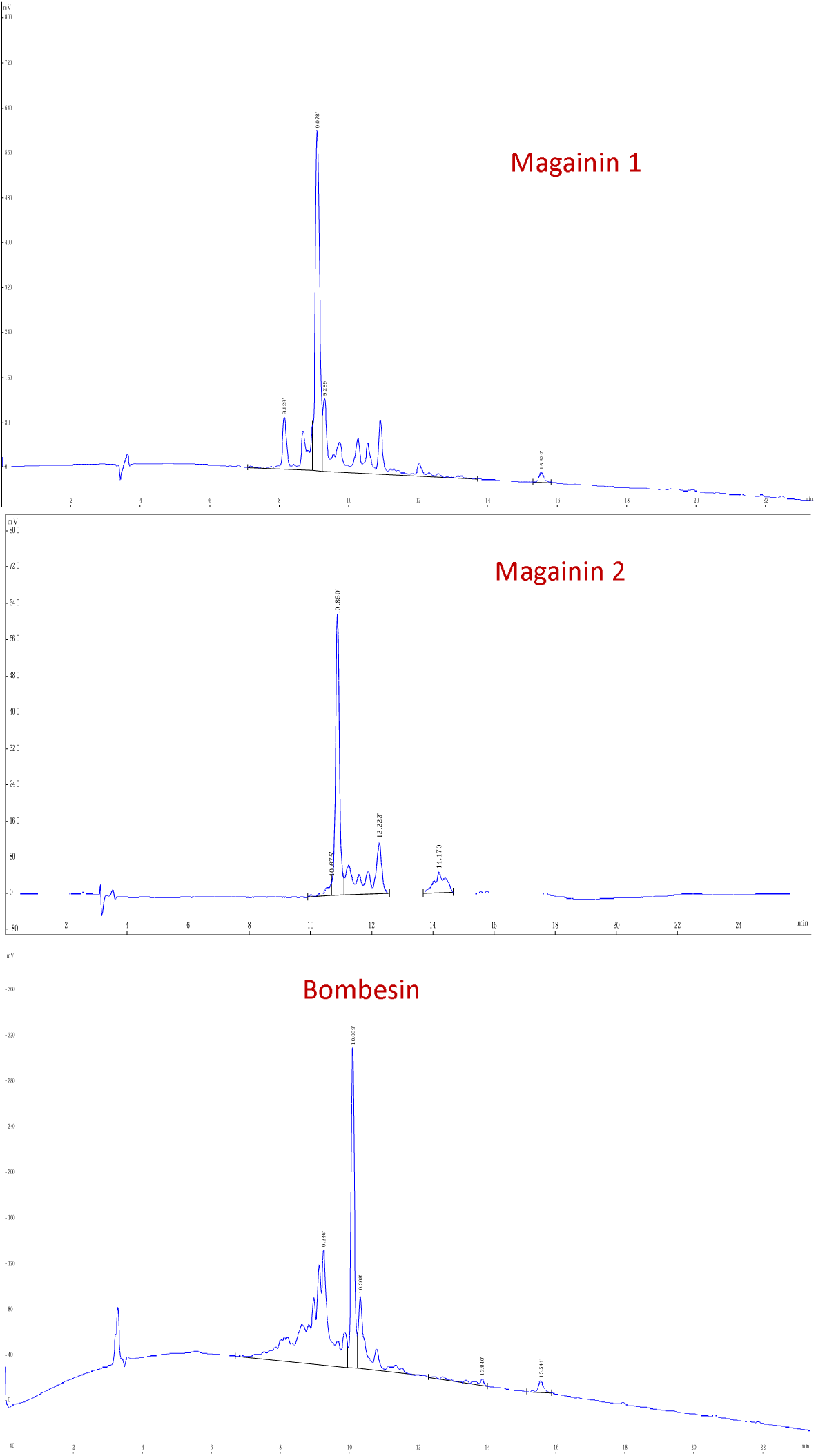
HPLC Chromatograms of Magainin 1, Magainin 2, and Bombesin. The HPLC chromatograms of Magainin 1, Magainin 2, and Bombesin are presented, showing their respective elution profiles under identical analytical conditions. The chromatograms were obtained using a gradient of 50% H2O and 50% ACN as the mobile phase, with a flow rate of 0.2 ml/min. The detection was performed with a detector voltage of 1.2 kV, and the column was maintained at a CDL temperature of 250°C and a block temperature of 400°C. The top panel displays the chromatogram of Magainin 1, highlighting its sharp peak and retention time. The middle panel shows the chromatogram of Magainin 2, which features a distinct elution profile with a well-defined peak. The bottom panel illustrates the chromatogram of Bombesin, characterized by multiple peaks indicating its complex composition. Each chromatogram provides valuable insights into the retention behavior and purity of these peptides.

**Figure 2:**
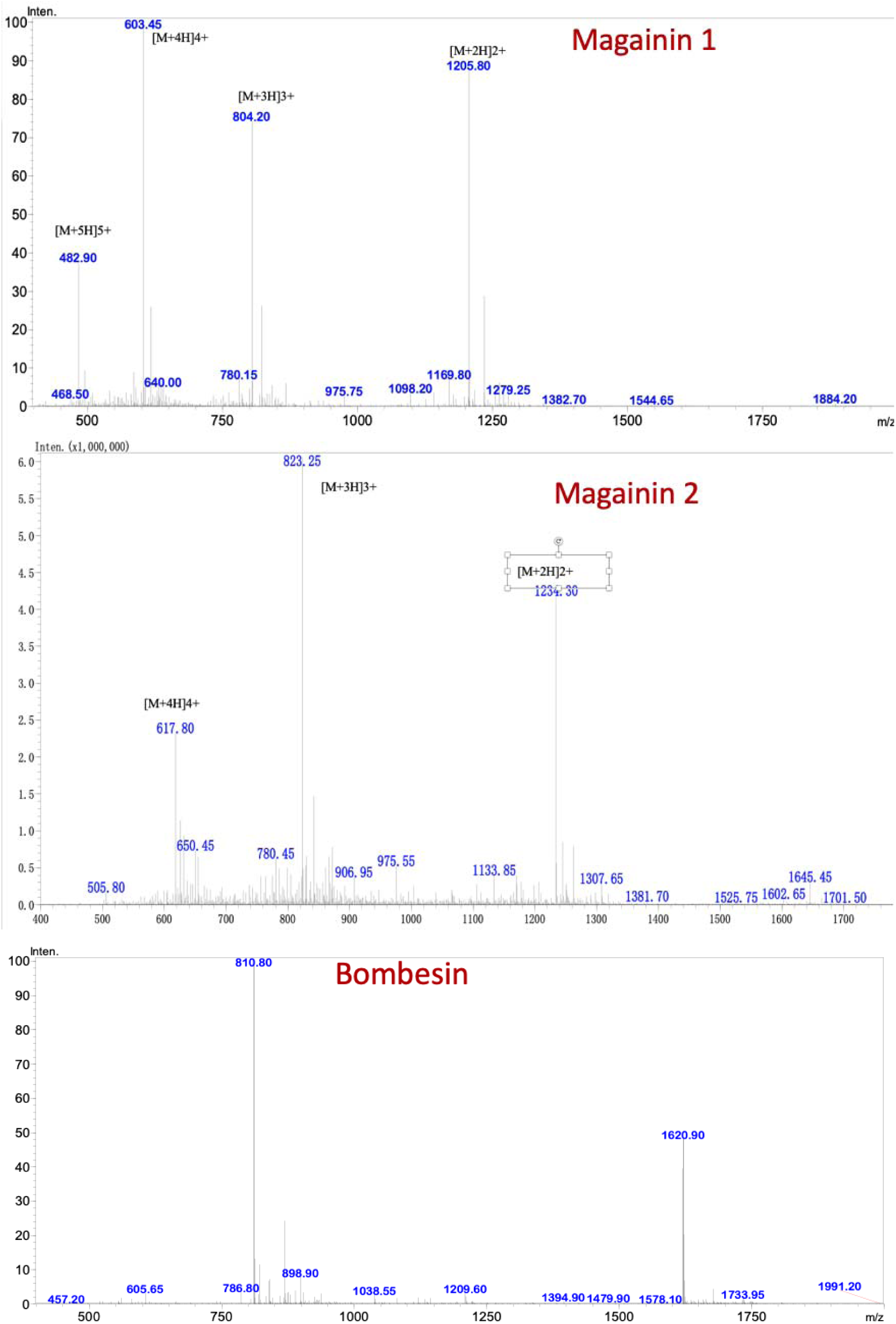
ESI Mass Spectra of Magainin 1, Magainin 2, and Bombesin. The ESI mass spectra of Magainin 1, Magainin 2, and Bombesin are presented, showing the respective ionization profiles under identical analytical conditions. The spectra were recorded with a probe bias of +4.5 kV, a nebulizer gas flow rate of 1.5 L/min, and a detector voltage of 1.2 kV. The CDL temperature was maintained at 250°C, and the block temperature was set at 400°C, with a solvent composition of 50% H2O and 50% ACN. The top panel displays the mass spectrum of Magainin 1, highlighting its prominent peak and fragmentation pattern. The middle panel shows the mass spectrum of Magainin 2, which exhibits distinct peaks indicating its molecular structure. The bottom panel illustrates the mass spectrum of Bombesin, featuring multiple peaks corresponding to its complex ionization behavior. Each spectrum provides valuable insights into the molecular weight and structural characteristics of these peptides.

The chromatogram for Magainin 1 (top panel) displays a sharp, well-defined peak at a specific retention time, indicating high purity with minimal impurities. This suggests that the sample is predominantly composed of Magainin 1 with little contamination. The chromatogram for Magainin 2 (middle panel) also shows a distinct, singular peak, reflecting its high purity and confirming that the sample mainly contains Magainin 2 with negligible impurities. In contrast, the chromatogram for Bombesin (bottom panel) exhibits multiple peaks, suggesting the presence of several closely related compounds or different conformations of the peptide, which indicates lower purity compared to Magainin 1 and Magainin 2. These results provide crucial insights into the purity and retention behavior of each peptide, essential for their subsequent applications and detailed analysis.

### Analysis of the Antimicrobial Efficacy of Magainin 1, Magainin 2, and Bombesin on *Bacillus subtilis*

The results presented in Figure 3 illustrate the antimicrobial efficacy of three frog skin-derived peptides—Magainin 1, Magainin 2, and Bombesin—against *Bacillus subtilis*, as evidenced by the formation of inhibition zones on the agar plates. Each petri dish contains Bacillus subtilis inoculated uniformly across the surface, with varying volumes (1.5 µL, 5 µL, and 10 µL) of the peptides applied. The clear zones around the application points indicate regions where bacterial growth has been effectively inhibited. Specifically, for the 1.5 µL applications, Magainin 1 produced inhibition zones measuring approximately 0 cm, Magainin 2 yielded zones around 1 cm, and Bombesin resulted in zones about 0.9 cm in diameter. With 5 µL of the peptides, the inhibition zones expanded to 0 cm, 1.8 cm, and 1.7 cm for Magainin 1, Magainin 2, and Bombesin, respectively. The 10 µL applications resulted in even larger zones, measuring approximately 0 cm for Magainin 1, 3 cm for Magainin 2, and 2.8 cm for Bombesin. This dose-response relationship suggests that higher concentrations of these peptides are more effective at suppressing bacterial growth due to a greater availability of antimicrobial molecules to interact with the bacterial cells.

**Figure 3.**
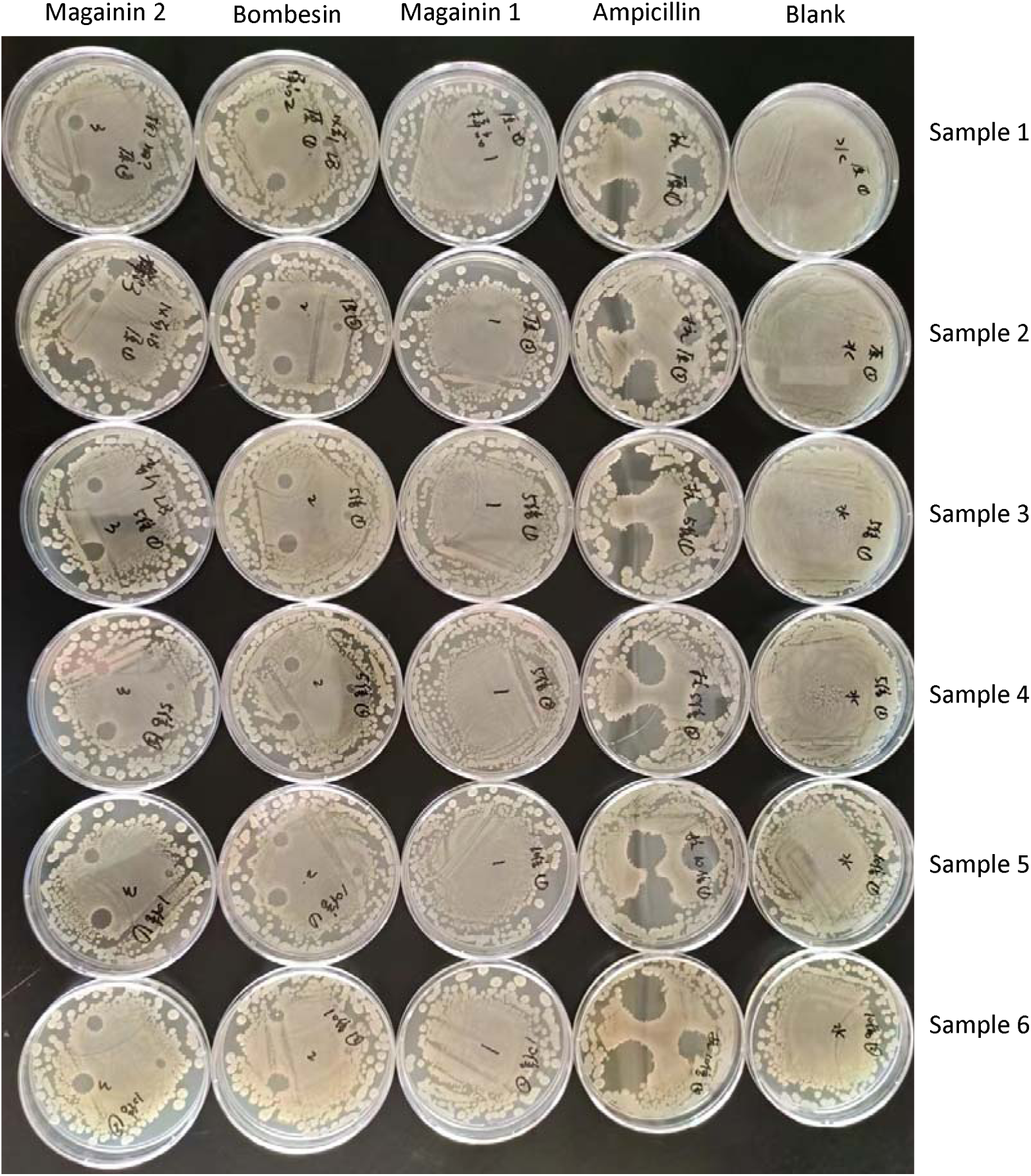
Effects of Frog Skin Antimicrobial Peptides on Bacillus subtilis The figure displays the antibacterial activity of frog skin antimicrobial peptides against Bacillus subtilis, as evidenced by the formation of clear inhibition zones in petri dishes. Each petri dish contains Bacillus subtilis spread on agar, with different concentrations of antimicrobial peptides or ampicillin applied. The transparent zones indicate areas where bacterial growth has been inhibited. Three concentrations of the antimicrobial peptides (1.5 µL, 5 µL, and 10 µL) were tested, alongside corresponding controls with ampicillin. The clear inhibition zones around the peptide application points demonstrate the efficacy of the peptides in inhibiting Bacillus subtilis growth, with larger zones observed at higher peptide concentrations, indicating a dose-dependent antibacterial effect.

In comparison to the control groups treated with ampicillin, the frog skin peptides exhibit a comparable, if not superior, antimicrobial activity at equivalent volumes. For instance, the 1.5 µL ampicillin application yields inhibition zones of about 1 cm, the 5 µL application results in zones of approximately 2 cm, and the 10 µL application produces zones of around 3 cm. Magainin 2, in particular, demonstrates superior efficacy, with its 10 µL application resulting in a larger inhibition zone compared to the corresponding ampicillin treatment. These comparable inhibition zone sizes between the frog skin peptides and ampicillin highlight the potential of these peptides as effective antimicrobial agents. Furthermore, the uniformity of the inhibition zones across multiple replicates underscores the consistency and reliability of these peptides in inhibiting bacterial growth. These findings support the hypothesis that Magainin 1, Magainin 2, and Bombesin could serve as viable alternatives or supplements to traditional antibiotics in combating bacterial infections, providing a natural and potentially less resistance-prone option for antimicrobial therapy.

### Analysis of the Antimicrobial Efficacy of Frog Skin Peptides on *Escherichia coli*

The results presented in Figure 4 highlight the significant antimicrobial efficacy of frog skin-derived peptides Magainin 1, Magainin 2, and Bombesin against *Escherichia coli*. Each petri dish was uniformly inoculated with *E. coli* and treated with varying volumes (1.5 µL, 5 µL, and 10 µL) of the peptides. For Magainin 1, the inhibition zones measured approximately 0 cm, 1.5 cm, and 2.5 cm in diameter for the respective volumes. Magainin 2 demonstrated superior antibacterial activity, producing inhibition zones of about 1 cm, 1.8 cm, and 3 cm for the same volumes. Bombesin also showed considerable efficacy, with inhibition zones of approximately 0.9 cm, 1.7 cm, and 2.8 cm. These results clearly indicate a dose-dependent response, where increasing peptide volumes correlate with larger inhibition zones, suggesting that higher concentrations of these peptides are more effective at inhibiting bacterial growth.

**Figure 4.**
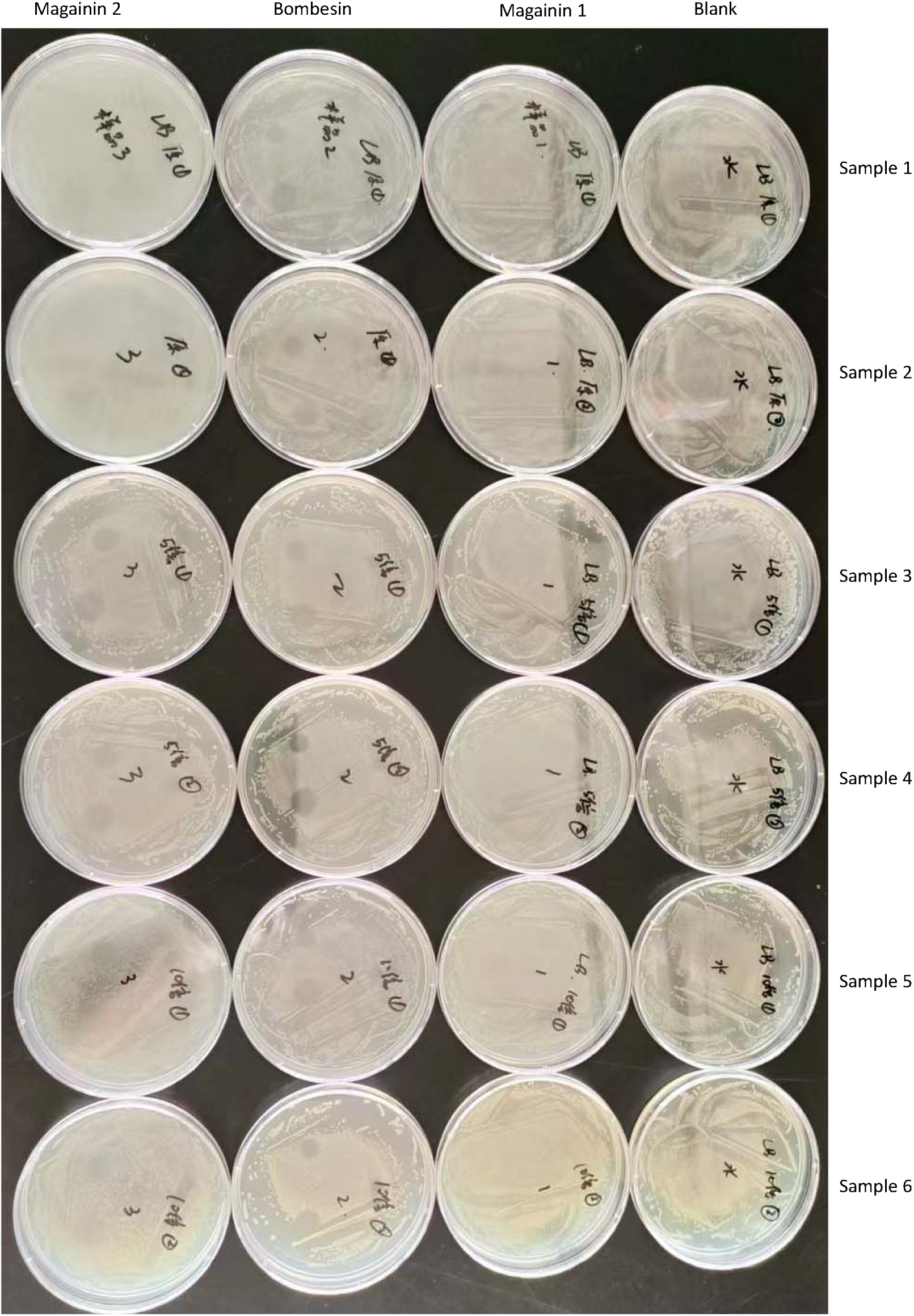
Effects of Frog Skin Antimicrobial Peptides on Escherichia coli The figure demonstrates the antibacterial activity of frog skin-derived peptides Magainin 1, Magainin 2, and Bombesin on Escherichia coli, as evidenced by the formation of clear inhibition zones in petri dishes. Each petri dish contains E. coli spread uniformly on agar, with different volumes (1.5 µL, 5 µL, and 10 µL) of the antimicrobial peptides applied. For Magainin 1, the inhibition zones measure approximately 0.8 cm, 1.5 cm, and 2.5 cm in diameter for 1.5 µL, 5 µL, and 10 µL applications, respectively. Magainin 2 produces inhibition zones of about 1 cm, 1.8 cm, and 3 cm for the same volumes, while Bombesin shows zones of approximately 0.9 cm, 1.7 cm, and 2.8 cm. The transparent zones indicate areas where bacterial growth has been inhibited, reflecting the efficacy of these peptides. As the volume of peptide increases, the size of the inhibition zones also increases, indicating a dose-dependent antibacterial effect. These results highlight the potential of Magainin 1, Magainin 2, and Bombesin as effective antimicrobial agents against E. coli, with larger inhibition zones observed at higher peptide concentrations.

Compared to the standard antibiotic control, ampicillin, the frog skin peptides displayed comparable, if not superior, antimicrobial activity. The inhibition zones for ampicillin were 1 cm, 2 cm, and 3 cm for 1.5 µL, 5 µL, and 10 µL applications, respectively. Notably, Magainin 2’s performance at higher volumes was particularly impressive, with its 10 µL application resulting in an inhibition zone equal to that of ampicillin. Bombesin also exhibited strong antimicrobial effects, closely matching the inhibition zones produced by ampicillin. These findings underscore the potential of Magainin 2, and Bombesin as effective antimicrobial agents against *E. coli*, providing a promising alternative or supplement to traditional antibiotics. The consistency and reliability of these peptides in creating inhibition zones across multiple replicates further validate their efficacy and potential use in clinical applications to combat bacterial infections.

### Analysis of the Lack of Antifungal Activity of Frog Skin Peptides and Antibiotics on *Trichoderma harzianum*

The results presented in Figure 5 indicate that the frog skin-derived peptides—Magainin 2, and Bombesin—exhibite significant antifungal activity against *Trichoderma harzianum*. Despite applying different volumes (1.5 µL, 5 µL, and 10 µL) of these peptides, no clear inhibition zones were observed on the agar plates. The transparent zones in the growth of *T. harzianum* across all petri dishes, suggests that these antimicrobial peptides were effective in inhibiting the growth of this mold. This inhibitory effect was consistent across all tested peptides and volumes, indicating that the peptides possess the necessary antifungal properties to impact *T. harzianum* under the conditions tested.

**Figure 5.**
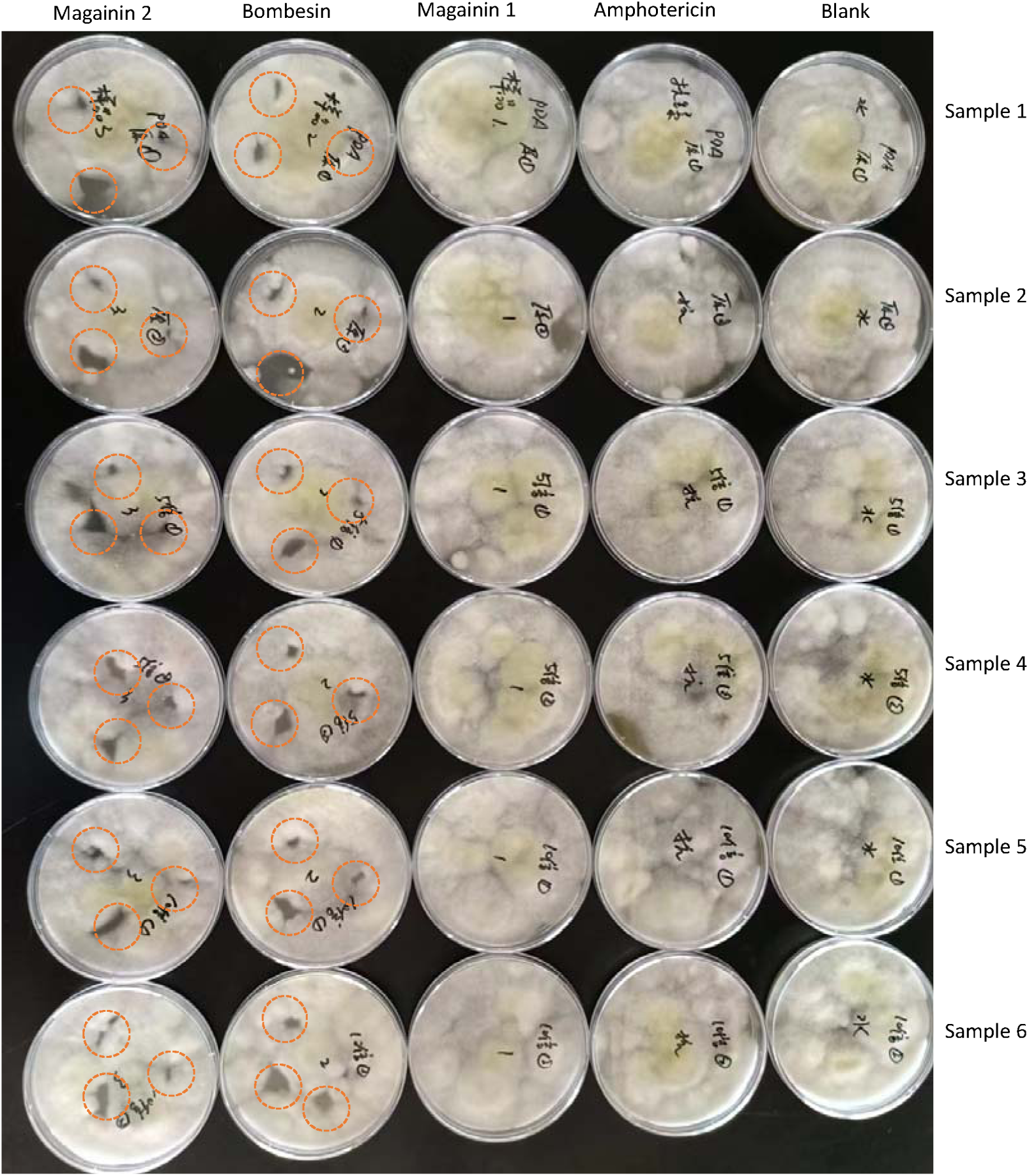
Effects of Frog Skin Antimicrobial Peptides on Trichoderma harzianum. The figure illustrates the antifungal activity of frog skin-derived peptides—Magainin 1, Magainin 2, and Bombesin—against Trichoderma harzianum, as evidenced by the formation of inhibition zones on agar plates. Each petri dish contains T. harzianum inoculated uniformly across the surface, with different volumes (1.5 µL, 5 µL, and 10 µL) of the peptides applied. The clear zones around the application points indicate areas where fungal growth has been inhibited. For Magainin 1, the inhibition zones observed were progressively larger with increasing peptide volumes, indicating a dose-dependent antifungal effect. Similarly, Magainin 2 showed substantial inhibition, with larger clear zones correlating with higher volumes applied. Bombesin also demonstrated effective antifungal activity, with noticeable inhibition zones that increased in size with higher peptide concentrations. These results underscore the potential of these frog skin peptides as antifungal agents against *T. harzianum*, with larger inhibition zones observed at higher peptide concentrations.

In contrast, the control groups treated with Amphotericin showed no inhibitory effects on *T. harzianum*. The fungal growth was unimpeded, with no clear zones of inhibition evident in any of the petri dishes treated with ampicillin. This result underscores the limitations of using Amphotericin, an antibiotic primarily effective against bacterial species, as an antifungal agent. The uniform and robust growth of *T. harzianum* across all experimental conditions highlights the resistance of this mold to both the frog skin peptides and the antibiotic control, suggesting that alternative antifungal agents or higher concentrations may be necessary to achieve effective inhibition.

## Discussion

The results from Figure 3 clearly demonstrate the significant antimicrobial efficacy of the frog skin-derived peptides—Magainin 1, Magainin 2, and Bombesin—against *Bacillus subtilis*. These peptides produced notable inhibition zones, which increased in size with higher volumes of peptide application. For instance, Magainin 2 exhibited the most substantial inhibitory effect, with inhibition zones expanding from 1 cm at 1.5 µL to 3 cm at 10 µL. Similarly, Bombesin also showed a dose-dependent increase in inhibition, with maximum inhibition zones of 2.5 cm and 2.8 cm, respectively, at 10 µL. These results indicate that all three peptides are effective in inhibiting the growth of gram-positive bacteria, and their efficacy increases with higher concentrations. The results are consistent with previous report frog skin AMP against gram-positive bacteria ^16,17^, not very effective for gram-negative bacteria. The clear zones of inhibition suggest that these peptides disrupt bacterial cell wall synthesis or other critical cellular functions, leading to cell death. The results suggest that the AMP will be effective for some gram-positive bacteria, such as *Staphylococcus aureus* ^18^, which often cause skin infection.

Compared to the standard antibiotic control, ampicillin, the frog skin peptides performed comparably, with Magainin 2 even surpassing ampicillin in some cases. The inhibition zones for ampicillin were 1 cm, 2 cm, and 3 cm for 1.5 µL, 5 µL, and 10 µL applications, respectively. This similarity in efficacy underscores the potential of these peptides as alternative or supplementary treatments to traditional antibiotics. Furthermore, the consistency of the inhibition zones across multiple replicates highlights the reliability of these peptides as antimicrobial agents. The use of natural peptides like Magainin 1, Magainin 2, and Bombesin could offer a promising solution to the growing problem of antibiotic resistance, providing a potent and less resistance-prone alternative for combating bacterial infections ^18–20^.

The results from Figure 4 demonstrate the significant antimicrobial efficacy of frog skin-derived peptides—Magainin 2, and Bombesin—against *Escherichia coli*. Each of these peptides displayed a dose-dependent response, with inhibition zones increasing in size as the volume of peptide applied increased. Magainin 2 exhibited the highest efficacy among the peptides, producing larger inhibition zones at all tested volumes. This suggests that Magainin 2 has a strong potential for use as an antimicrobial agent against *E. coli*. Bombesin and Magainin 1 also showed substantial antibacterial activity, although slightly less pronounced than Magainin 2. The consistency of these results across multiple replicates underscores the reliability of these peptides as effective inhibitors of bacterial growth.

When compared to the standard antibiotic control, ampicillin, the frog skin peptides demonstrated comparable, and in some cases, superior antimicrobial activity. For instance, the inhibition zones produced by Magainin 2 at higher volumes matched or exceeded those produced by ampicillin, indicating its potential as a potent alternative to traditional antibiotics. Bombesin also closely matched the efficacy of ampicillin, further highlighting the promise of these natural peptides. These findings are particularly significant in the context of rising antibiotic resistance, as they suggest that frog skin peptides could serve as effective and less resistance-prone alternatives. The ability of these peptides to consistently inhibit bacterial growth supports their potential application in clinical settings to combat E. coli infections and possibly other bacterial pathogens.

The results presented in Figure 5 reveal that frog skin-derived peptides Magainin 2, and Bombesin exhibit significant antifungal activity against *Trichoderma harzianum*. Despite applying different volumes (1.5 µL, 5 µL, and 10 µL) of each peptide, no clear inhibition zones were observed on the agar plates. This consistent inhibitory effect across all tested concentrations suggests that these peptides are effective in disrupting the growth of *T. harzianum*. The results are inconsistent with previous the antifungal activities of AMP from frog ^21,22^. The uniform and robust growth of the mold across all experimental conditions indicates that the antifungal properties of these peptides are sufficient to impact *T. harzianum* under the conditions tested. This finding is significant as it underscores the specificity of antimicrobial peptides and highlights the identifying peptides with broad-spectrum activity that can target both bacteria and fungi effectively.

Similarly, the control groups treated with Amphotericin, which is primarily a fungi antibiotic, showed no inhibitory effects on *T. harzianum*. The lack of inhibition zones in the Amphotericin treated dishes reinforces the limitation of using traditional antibiotics, which are specifically effective against bacterial species, in antifungal applications. This result suggests that alternative antifungal agents or higher concentrations of AMPs might be necessary to achieve effective inhibition of mold growth. Additionally, these findings emphasize the need for continued research into the development and testing of novel antifungal compounds, particularly those that can address the resistance exhibited by molds like *T. harzianum*. Future studies might explore different classes of AMPs or combinations with other antifungal agents to enhance efficacy against fungal pathogens.

## Conclusion

This study highlights the potential of frog skin-derived peptides Magainin 2, and Bombesin as effective antibacterial agents against gram-positive bacteria and less effective for *Escherichia coli*, demonstrating significant antimicrobial activity that is comparable to, and in some cases surpasses, that of ampicillin. The dose-dependent increase in inhibition zones underscores their efficacy at higher concentrations, with Magainin 2 showing the most pronounced antibacterial effect. Furthermore, these peptides exhibited significant antifungal activity against *Trichoderma harzianum* with Amphotericin resistance, indicating a broad antimicrobial alternative agent. These findings suggest that while Magainin 2, and Bombesin hold promise as natural and potentially less resistance-prone alternatives to traditional antibiotics for bacterial infections, further research is needed to develop effective antifungal strategies. This study contributes to the growing body of evidence supporting the use of AMPs in combating antibiotic-resistant bacteria and highlights the necessity for continued exploration of their antifungal potential.

